# A synthetic likelihood solution to the silent synapse estimation problem

**DOI:** 10.1101/781898

**Authors:** Michael Lynn, Kevin F.H. Lee, Cary Soares, Richard Naud, Jean-Claude Béïque

## Abstract

Functional features of populations of synapses are typically inferred from random electrophysiological sampling of small subsets of synapses. Are these samples unbiased? Here, we developed a biophysically constrained statistical framework for addressing this question and applied it to assess the performance of a widely used method based on a failure-rate analysis to quantify the occurrence of silent (AMPAR-lacking) synapses in neural networks. We simulated this method in silico and found that it is characterized by strong and systematic biases, poor reliability and weak statistical power. Key conclusions were validated by whole-cell recordings from hippocampal neurons. To address these shortcomings, we developed a simulator of the experimental protocol and used it to compute a synthetic likelihood. By maximizing the likelihood, we inferred silent synapse fraction with no bias, low variance and superior statistical power over alternatives. Together, this generalizable approach highlights how a simulator of experimental methodologies can substantially improve the estimation of physiological properties.

## Introduction

Activity-dependent synaptic plasticity has captured global research attention for well over four decades due to its hypothesized role in learning and memory (1). Silent synapses contain only NMDA receptors, not AMPA receptors, and represent preferential sites for receptor insertion during long-term potentiation (LTP; 2, 3). The insertion of AMPARs is generally accompanied by an increase in spine size (structural plasticity; 29, 47-49), linking structure and function (see 28, 50 for mechanistic details). The relative proportion of silent synapses in neural circuits is believed to be a fundamental determinant of the plasticity potential of neural networks. Silent synapses are more prevalent in developing networks (4, 5) yet continue to play a role in adult brain circuits for instance in newborn neurons (6), and mature synapses can at times undergo ‘silencing’ by removal of AMPARs (7). Furthermore, addictive drugs such as cocaine and morphine have been suggested to induce *de novo* formation of silent synapse in reward-related pathways of adult animals (8-11). More generally, theoretical work has shown that high fractions of ‘potential’ synapses (*e*.*g*., silent) are associated with optimal information storage over a range of conditions (12-14). Together, whereas these observations point to the fundamental role of silent synapses in both developing and mature brain circuits, they also emphasize the corollary challenge of quantifying with precision their occurrence in synaptic populations with experimentally realistic throughput.

While sound methods exist to demonstrate the presence of individual silent synapses (Fig. 1Ai-ii) (2, 3), these binary single-synapse classification methods can only estimate the fraction of silent synapses in a population by pooling across repeated single-synapse interrogations either using a minimum stimulation paradigm (4, 5), or glutamate uncaging techniques (22, 29). In contrast, multi-synapse classification methods, such as the failure-rate analysis (FRA) method, have been developed to provide an estimate of the fraction of silent synapses in a larger synaptic population for every experimental attempt: here an experimenter records from a small, but unknown, number of synapses, and the FRA formalism estimates the proportion of these which are silent by means of a comparison of failure rates (Fig. 1Bi-ii) (methodology reviewed in 15). A number of studies have used this method to quantify changes in silent synapse fraction, notably in rodent models of drug addiction, but also during critical periods of plasticity in the visual cortex and hippocampus (8-11; 16-21, 32, 45). Yet, although FRA seeks to scale up the throughput of silent synapse estimation, its scalability hinges on its reliability and accuracy (especially compared with traditional binary classification methods), which have not been investigated with targeted experimental and statistical analyses.

**Figure 1:**
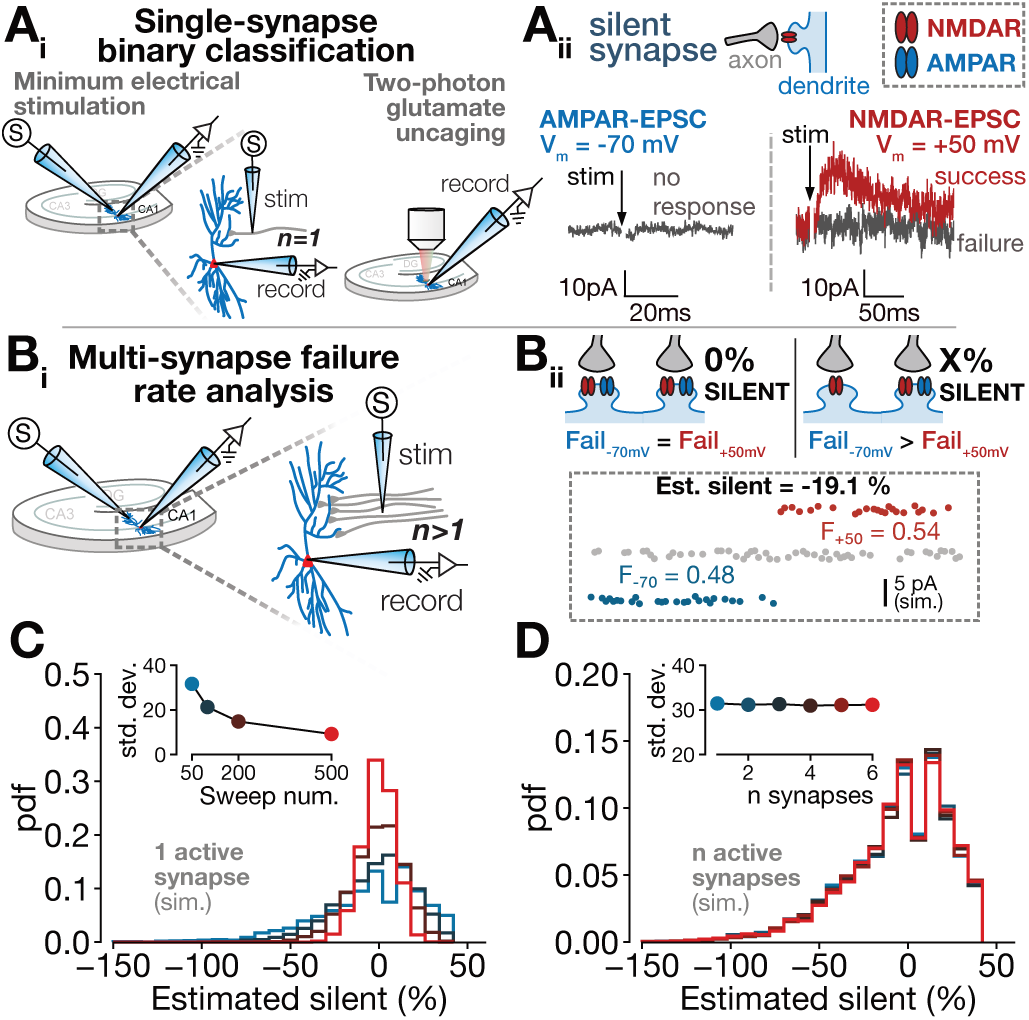
The FRA technique returns high-variance estimates under conditions with no silent synapses present. **Ai:** Single-synapse classification techniques rely on electrical (left) or optical (right) stimulation of a single synapse. **Aii**: The responses (here averaged electrical traces), are classified as successes and failures at -70mV (left) and 50mV (right). **Bi:** In contrast, the multi-synapse failure rate analysis method relies on stimulating multiple synapses and (**Bii**) comparing failure rates at two potentials to estimate silent fraction (inset shows example calculation). **C**: FRA estimate distributions for a single active neuron, as the number of sweeps per voltage condition is varied (inset, colors indicate sweep num.). **D:** FRA estimate distributions for varying numbers of active synapses (inset, colors indicate synapse num.), with mean failure rate held constant at 0.5.

Here, we first employ a combination of biophysically constrained numerical simulations and electrophysiological recordings to examine the performance of existing methods for silent synapse detection and quantification. Our analysis reveals substantial and general limits on the reliability and validity of the silent synapse fraction estimates obtained using the FRA formalism. To overcome these inherent limitations, we build on approaches developed in the dynamical systems field and employed our computational model to generate an approximate maximum-likelihood estimator. This approach provides dramatic improvements in estimator bias and variance. A power analysis reveals that the enhanced estimator can resolve fine-grained changes in silent fraction with experimentally feasible sample sizes. Our estimator does not require a novel set of observations and can thus be deployed on existing FRA datasets. These tools and findings make large scale interrogation of changes in silent synapses possible and reliable.

## Results

While minimum stimulation experiments have been critically important for demonstrating the existence of silent synapses, they do not formally allow estimation of the true silent synapse fraction (*s*_*t*_) contained in a synaptic population. A simple experimental technique, the failure rate analysis (FRA), has been developed to provide a quantitative estimate (*ŝ*_*FRA*_) of the true underlying silent fraction, *s*_*t*_. In this method, the rate of synaptic failures at two holding potentials (−60 mV and +40 mV) are used as inputs to a mathematical equation (see STAR Methods for derivation), which returns an estimated fraction of silent synapses. Despite this formalism, it is unclear how biophysical variables such as stochastic neurotransmitter release and the inherent variability in the number of synapses stimulated between experiments shape the statistical power and discriminability of this method. Here, we use a set of increasingly constrained and experimentally realistic numerical simulations and statistical analysis to examine the performance and general usefulness of this estimator.

### A. Nonsilent synaptic populations and FRA estimator performance

As a first step, we directly determined the extent to which the stochasticity of synaptic release affect the FRA estimator. We thus numerically simulated the simplest scenario where a single, non-silent synapse with probability of release *Pr* = 0.5 is electrically stimulated (Fig. 1Bii, inset). Henceforth we will refer to non-silent synapses as active synapses (i.e., synapses containing both AMPA and NMDARs). We simulated Bernoulli-distributed trials (failures and successes) in 50-sweep epochs. The FRA technique, applied to a single, active synapse (i.e., *s*_*t*_ = 0) should return *ŝ*_*FRA*_ ≈ 0. With 50 sweeps, the distribution of *ŝ*_*FRA*_ across multiple experiments (Fig. 1C, blue) was strikingly broad (S.D. = 31.3%), non-Gaussian, had high kurtosis (kurtosis = 3.00) and a prominent left tail (skewness = -1.01). These attributes therefore make the FRA estimator unsuitable for the traditionally-used parametric tests that assume normality.

Furthermore, 45.3% of the *ŝ*_*FRA*_ distribution falls below zero, which are biologically irrelevant estimates. Since negative silent synapse values are conspicuously absent from published datasets using FRA (8-11, 16-21), it is possible that these negative values returned from the equation were arbitrarily set to 0% silent (i.e., systematically ‘zeroed’). This procedure would artificially inflate the mean silent synapse estimate.

We next asked whether the FRA estimator’s accuracy could be improved by simply increasing the number of sweeps from the recommended value of 50 (15) (Fig. 1C). Increasing the number of sweeps per epoch to 500 still produced a remarkably variable estimate (Fig. 1C inset; 500 sweeps: S.D. = 9.1%).

Alternatively, recordings from multiple synapses may improve the reliability of *ŝ*_*FRA*_, due to the intuitive averaging out of stochastic binary events. We thus numerically simulated FRA experiments on small numbers of synapses, first analytically calculating a fixed Pr for all synapses such that the mean failure rate was 50% (Fig. 1D). Surprisingly, *ŝ*_*FRA*_ variability was virtually identical over the entire range of active synapse numbers (Fig. 1D, inset). Therefore, in the simple case of non silent populations, the variability of the FRA estimates is not markedly improved when monitoring multiple synapses.

We considered the generalizability of our findings by asking how *ŝ*_*FRA*_ estimates varied over a wide Pr range and ensemble synapse number (Fig. S1). Our analysis revealed that the FRA estimator variance, surprisingly, is inversely proportional to synapse determinism (i.e., as *Pr* → 0^+^ and as *Pr* → 1^−^) and does not significantly decrease with any combination of Pr and synapse number. Taken together, these findings outline the general inability of the FRA method to return reliable (i.e., low variance) estimates of silent synapse fraction in simple conditions with no silent synapses present (*s*_*t*_ = 0%).

### B. Silent synapse-containing synaptic populations and FRA estimator performance

Under conditions where silent synapses are present (*s*_*t*_ > 0%), the quantitative accuracy of the FRA method is naturally dependent on the particular arrangement of synapses sampled. It is not intuitively obvious whether traditional electrical stimulation selects an unbiased subsample of a larger synaptic population, either in terms of silent fraction or of individual synapse properties such as release probability. Therefore, before assessing the accuracy of the FRA equation, we first developed an obligatory intermediate method to simulate experimentally feasible ‘draws’ of synapse sets, with varying release probabilities, from larger mixed populations (Fig. 2A).

**Figure 2:**
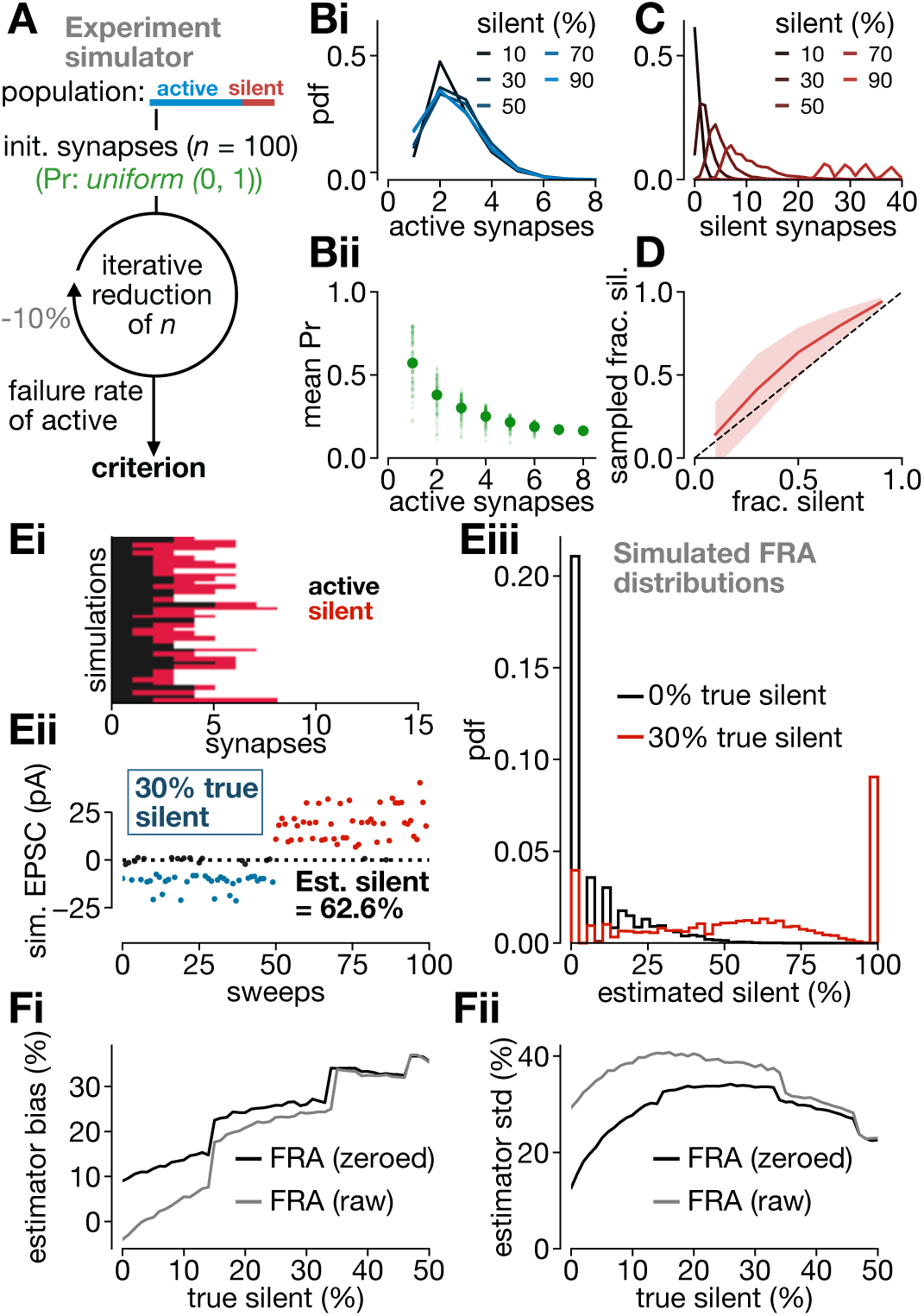
Experimentally bounded simulations reveal a fundamental bias inherent in bulk electrical stimulation. **A:** Schematic depicting experimental simulator approach. **Bi:** Probability distribution of number of active synapses obtained during each simulation. **Bii:** Release probability of active synapses. Small dots denote release probability for each simulation, while large dots denote the mean across simulations. **C:** Probability distribution of number of silent synapses obtained during each simulation. **D:** Sampled fraction silent is shown against the true fraction silent. **Ei**: Depiction of returned samples of silent/active synapses. **Eii**: Depiction of a simulated FRA experiment performed on a sample of synapses returned from the iterative reduction. **Eiii**: Distribution of FRA estimates for two true silent values. **Fi:** FRA estimator bias for non-zeroed and zeroed distributions. **Fii**: FRA estimator standard deviation for non-zeroed and zeroed distributions.

Briefly, we started with initial sets of 100 synapses with some true fraction silent (*s*_*t*_). Release probabilities were uniformly distributed (*Pr* ∼ *U*(0,1)). We next simulated as faithfully as possible a typical FRA electrophysiological experiment. Thus, the initial set of activated synapses was subjected to multiple rounds of stochastic synapse elimination (10% per round) corresponding to a decrease in electrical stimulation intensity that an experimenter typically performs during an FRA experiment. The rounds of elimination were ended when the failure rate of active synapses reached between 20% and 80%, as dictated by described experimental guidelines (15). Synapse sets were discarded if they did not reach this criterion before the elimination of active synapses. The simulation thus returned stochastic, small subpopulations of silent and active synapses, each with an associated release probability, and with the failure rate range as the main constraint.

We found that the number of active synapses in each subpopulation reaching the selection criterion was very small - typically between 1 and 5 (Fig. 2Bi). As one may intuit, large numbers of synapses in a draw are rare because they require very low *Pr* for every synapse to achieve the desired failure rate (Fig. 2Bii). This experimentally-derived selection criterion therefore leads to a bias towards sampling synapses with low Pr. The number of silent synapses in the draws scaled with *s*_*t*_, as expected (Fig. 2C). However, the sampled subsets were consistently enriched in silent synapses compared to the ground-truth population (Fig. 2D). This intriguing bias arises from the simple fact that the failure-rate criterion *de facto* solely depends on active synapse monitoring: therefore, the last elimination round ineluctably eliminates an active, rather than a silent, synapse. On average, the number of active synapses is therefore under-represented in the sample, which leads to a quasi-systematic overestimation of the silent fraction. Given that the small size of the sampled synaptic population (1-5), this systematic bias is consequential. This appears to be a general feature of the FRA method when stimulation amplitude is adjusted to reach a particular failure rate.

We next considered the performance of the FRA estimator under this experimentally realistic scenario.

Individual simulations returned highly variable synapse sets (Fig. 2Ei) and inaccurate estimates (*ŝ*_*FRA*_) (Fig. 2Eii). Over multiple simulated experiments, the estimated silent synapse fraction showed distinctly non-normal distributions (Fig. 2Eiii). We next undertook a formal analysis of the FRA estimator’s bias and variance. The bias of *ŝ*_*FRA*_ increased with the true silent fraction, *s*_*t*_, and this occurred irrespective of whether negative estimates were erroneously set to zero (‘zeroing’) or not (Fig. 2Fi). The standard deviation of the estimate was high across all silent fractions (Fig. 2Fii), reaching a maximum of 40% with zeroing applied. We conclude that the FRA estimator displays a high positive bias and large variance, and thus is a flawed estimation technique.

While we drew the release probability of individual synapses from a uniform distribution following previous work assessing Pr from iGluSnFR-mediated single-synapse release events (22), we additionally asked whether different release probability distributions of a synaptic population would influence the FRA estimator’s performance. First, we considered the gamma distribution (parameters: *λ* = 5.8; *n* = 3) reported from staining-destaining experiments (Fig. 1G in ref. 23). This distribution produced negligible changes in both the synapse selection bias, and in the bias and variance of the estimator (Fig. S2). We additionally simulated the case with extremely small release probabilities (*λ* = 5.8, *n* = 1) (Fig. S3). Although the mean number of active synapses rises from ∼3 to ∼10 (Fig. S3Bi), the estimator with zeroing applied still shows comparable bias (27% bias at 50% silent, compared to 35% bias at 50% silent for a uniform distribution) and variance (peak standard deviation of 30%) (Fig. S3Fi-ii). These results demonstrate the significant propensity of the FRA estimator towards large positive biases and large variances over a wide range of possible synaptic release probability distributions.

### C. Electrophysiological verification of numerically simulated FRA estimate inaccuracies

We next sought to gain an experimental validation of the inherent propensity of the FRA approach to return variable estimates of silent synapse fraction. We thus performed voltage-clamp recordings from CA1 neurons from p14-15 mice while electrically stimulating Schaffer’s collateral synapses (Fig. 3A; see STAR Methods). As per recommended practice for typical FRA experiments, we tuned the stimulation intensity such that we obtained clear monosynaptic currents with ∼50% failure rates (Fig. 3B,C). We performed FRA experiments in triplicate for each recording from individual CA1 neurons (Fig. 3D), and we verified that the evoked EPSC amplitudes at both -70mV and +40mV were stable between these successive iterations (Fig. 3Eii), confirming that we were stimulating the same subsets of synapses during each FRA iteration.

**Figure 3:**
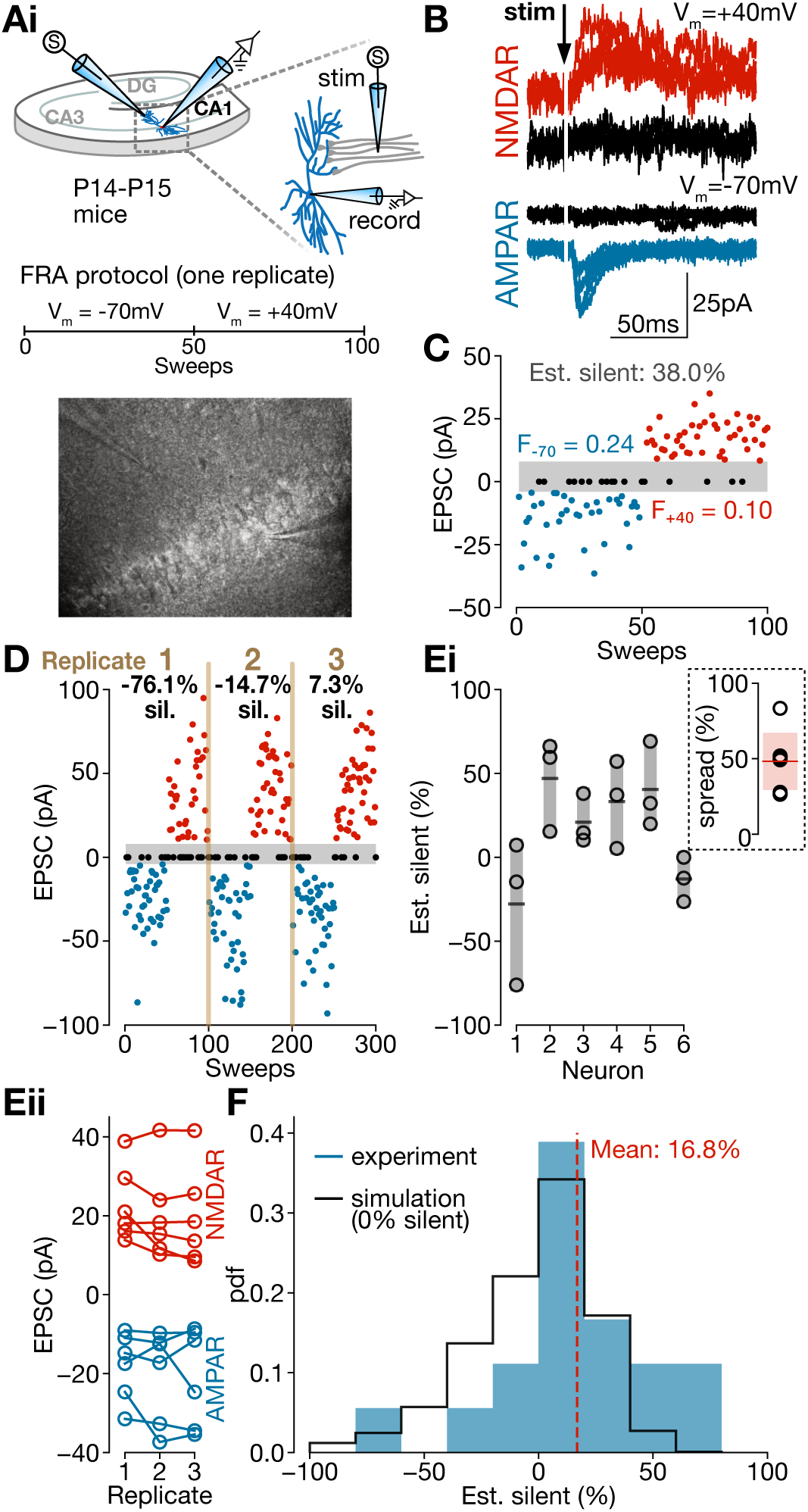
Electrophysiological recordings at CA3-CA1 synapses are consistent with simulated FRA distributions. **A:** Experimental schematic illustrating recording from CA1 neurons while stimulating CA3 afferents. **B:** Representative traces showing successes and failures at Vm = +40mV (top) and Vm = -70mV (bottom). **C:** Exemplar experiment showing failure rates and estimated silent fraction. **D:** Three replicates of an FRA experiment on a single neuron returns highly variable estimates consistent with biophysical limitations of the FRA method. **Ei:** Replicates for each neuron (circles) and the mean of these estimates (bar). Inset depicts estimate spread for each neuron. **Eii**: EPSC amplitude stability across replicates for each neuron. **F:** Histogram of all replicates (blue) overlaid with simulated histogram from null distribution (0% silent, black). Distributions are significantly different (p = 0.049, Kolmogorov-Smirnov two-sided test).

Since we performed the FRA experiments in triplicate for each recordings, we computed three independent *ŝ*_*FRA*_ estimates of the same synaptic population. As predicted from our simulations, these estimates were highly variable within the same neuron (Fig. 3Ei), and often returned biologically irrelevant negative values (Fig. 3F). Because we cannot directly measure *s*_*t*_, we are unable to formally compare the bias and variance of the estimator with experimental ground truth, but these results nonetheless demonstrate the strong agreement between our computational model and experimental findings in documenting the fundamental limits of the FRA formalism.

### D. An improved estimation technique based on synthetic likelihood functions

In principle, it should nonetheless be possible to extract reliable information about the true state of a synaptic population from unreliable and biased estimates returned by the FRA method. For instance, if we knew the distributions of failure rates at both hyperpolarized and depolarized potentials for each *s*_*t*_, we could write expressions for likelihood functions and construct a maximum-likelihood estimator (MLE). The logic of this approach is to first determine the analytical function establishing the probability of an observation given some model parameters, and then to find the model parameter (here the silent synapse fraction) that maximizes this likelihood function. This type of estimator is of particular interest here since MLE minimizes the variance, at least asymptotically (37).

In the present case, however, the likelihood functions are analytically intractable, as they result from a complex combination of the sampling protocol and the stochastic properties of synapses. Some simplifying assumptions could be made to reach a closed-form likelihood function, but this would require assuming that release probability distributions are independent of sampled synapse number, and that the sampled subsets themselves reflect unbiased samples from the population (both assumptions are shown by our model to be false). Alternatively, popular approaches for noisy dynamical systems sidestep analytical solutions and rather construct synthetic likelihood functions from Monte Carlo simulations (33-35). Here, the dynamical system is iteratively simulated and summary statistics are extracted, allowing to likelihood-based estimation from a stochastic model. These likelihood function are approximations only in the sense that they are based on a finite number of stochastic simulations, but will converge to the true likelihood function. We use this formalism to construct an estimator, FRA-MLE, from synthetic likelihood functions (Fig. S4A-B) where the summary statistic is the biased FRA estimate itself (*ŝ*_*FRA*_) (See STAR Methods for more details on why this summary statistic was chosen). Since our noisy sampling model already incorporates technical choices made by experimenters (e.g. the range of acceptable failure rates), this allows us the flexibility to construct estimators, *ŝ*_*FRA*−*MLE*_(***θ***) which are a function of an experimental parameter vector ***θ*** containing methodological variables and synaptic properties.

In contrast with the large bias found for the FRA estimator, we found that *ŝ*_*FRA*−*MLE*_ is entirely unbiased (Fig. 4A, Fig. 2Fi). Also in contrast with the large variance found for the FRA estimator (Fig 2Fii), we found that although the variance of *ŝ*_*FRA*−*MLE*_ increases with ground truth silent fraction, it remains lower than that of *ŝ*_*FRA*_ (Fig. 4B; Fig. S4C). Therefore, the new FRA-MLE method circumvents the inherent sampling bias and high variance of the original *ŝ*_*FRA*_.

**Figure 4:**
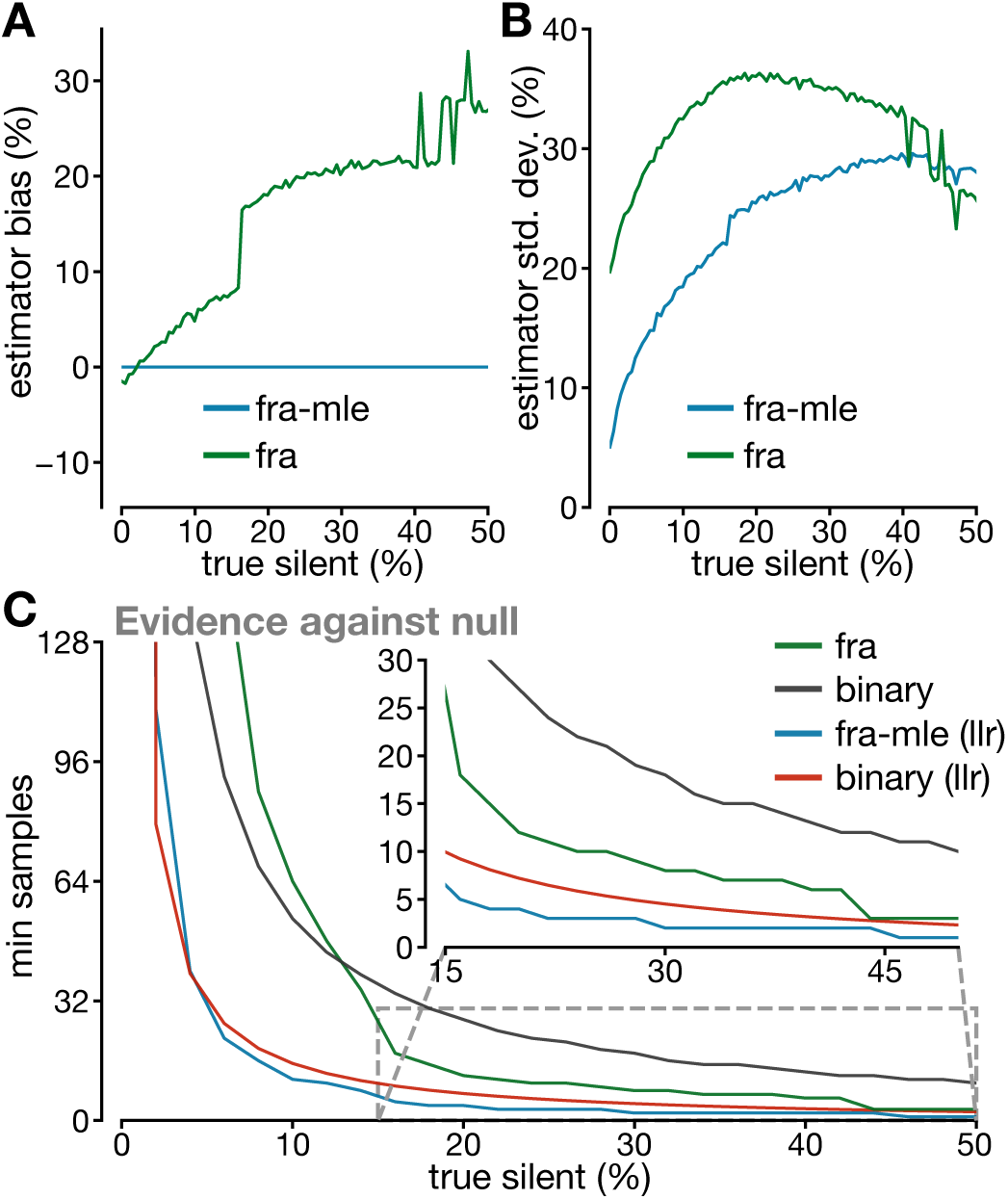
Statistical properties of FRA-MLE estimator compared with other methods. Shown are FRA -MLE (blue) and FRA (green) estimator bias (**A**) and standard deviation (**B**). **C:** Minimum samples required for each estimator to discriminate true silent fractions from a null population where no silent silent synapses are present. Inset depicts a zoomed view of 15-50% silent (*α* = 0.05, *β* = 0.2 for all cases).

As a final test, we applied FRA-MLE to the experimental values collected from hippocampus (Fig. S4D-F). We used FRA-MLE to arrive at a maximally likely estimate, *ŝ*_*FRA*−*MLE*_, across all experimental observations. We found that the joint likelihood function was sharply tuned, peaking at a maximal likelihood estimate of 11.4% silent for our experiments (Fig. S4E). This was substantially lower than the estimate obtained by simply taking the mean of all experimental FRA values (16.8%), in agreement with our findings of a pervasive positive bias for FRA. We additionally verified our estimator and numerical simulation approach as follows: We employed our statistical sampling model (*i*.*e*., successive rounds of synapse elimination) to construct a hypothetical experimental distribution of FRA estimates for the maximally likely silent fraction *ŝ*_*FRA*−*MLE*_ = 11.4% obtained for our experiments. We found that this model-generated distribution was not significantly different from the experimental distribution (Fig. S4F; p = 0.99; KS-test, two-tailed). Together, these findings reiterate the general propensity of FRA to return positively biased estimates, and demonstrate the ability of FRA-MLE to extract not just locally optimal solutions, but solutions which are also in good quantitative correspondence with experimental distributions.

### E. A power analysis quantifies performance improvements of the FRA-MLE estimator

To achieve a high-throughput characterization, one needs to take two sets of *n* samples from either of two separate conditions, and conclude with an experimentally realistic *n* whether the two conditions do not have the same proportion of silent synapses (*i*.*e*. reject the null hypothesis). We used the framework of power analyses to ask whether our FRA-MLE estimator compares favorably with other methods, classic FRA as well as single-synapse binary classification methods, including minimum stimulation experiments as well as more modern 2-photon glutamate uncaging methods. We calculated the minimum sample size, *n*_*min*_, required to discriminate silent-containing populations from an active-only population utilizing each technique. Depending on the estimator, calculating *n*_*min*_ is done by either analytically or numerically calculating the minimum sample size to achieve a threshold statistical power (*i*.*e*. a threshold false negative rate; see STAR Methods). More specifically, since the FRA-MLE method calculates an approximate likelihood function explicitly, we could use the log-likelihood ratio (LLR) to test against our null hypothesis (38), an approach that is not possible with the FRA estimator. We can compare against the binary method (Fig. 1A) either using a LLR as done for the FRA-MLE estimator or with a paired sample test as done with the FRA method. Thus we compare four statistical protocols: FRA estimates using a paired test, FRA-MLE estimates using LLR, binary classification using a paired test and binary classification using LLR.

Figure 4C shows *n*_*min*_ against the true silent synapse fraction. As is expected, we found that more samples are required to discriminate smaller differences in silent synapse fractions from a fully active synapse population. Importantly, we found a dramatic improvement in statistical power afforded by the FRA-MLE technique (Fig. 4C) compared to FRA. While the FRA estimator required *n*_*min*_ = 36 to discriminate populations containing less than 15% silent, FRA-MLE required only *n*_*min*_ = 8. Averaged across all silent fractions, FRA-MLE achieved an identical statistical power to the FRA estimator with a 3.82-fold reduction in sample size. Binary classification required 1.88-fold more samples than FRA, although when LLR-based hypothesis testing was incorporated, the binary technique now required fewer samples than FRA such that its power almost matched that of the FRA-MLE method. Together, this power analysis shows that the FRA-MLE estimator renders more fine-grained silent fractions detectable with fewer samples than comparable methods.

## Discussion

It has been long believed that the plasticity potential of a network is in part determined by the fraction of synapses which exist in a silent state, and the proportion of silent synapses is regulated both throughout development (4-5) and in response to diseases or drugs (8-20). Despite a growing appreciation of the overall importance of silent synapses in mature neural networks, a systematic analysis of quantification methods for this parameter is, to our knowledge, lacking. Here, using statistical inference methods alongside electrophysiological recordings, we have demonstrated the significant bias and variance inherent to the popular failure-rate analysis method of silent synapse quantification. Through a set of numerically simulated likelihood functions relying on minimal assumptions, we have furthermore proposed and validated an alternate maximum-likelihood estimator which corrects for these flaws. This approach illustrates the advantages of a model-based statistical inference approach.

One assumption of our approach is that release probabilities across a synaptic population are drawn from a uniform distribution (22). It is possible that specific constellations of central synapses in the brain may deviate from the later assumption. For instance, a population dominated by very low release probability synapses would inevitably result in the sampling of a greater number of synapses, which could in principle increase the statistical power of the method and potentially decrease bias. To address this possibility, we considered the release probabilities derived from staining-destaining experiments in hippocampus that did not follow a uniform distribution (Fig. S2; ref. 23) and found that it induced minimal changes in the mean estimates of sampled synapses. These considerations in effect highlight a strength of the MLE model-based approach in that experimenters can ensure that the statistical model is precisely calibrated to their experiment’s methodology (e.g., here, to a specified release probability distribution).

We also note that any estimation of the failure rate itself is dependent on correctly binarizing a set of noisy analog traces into synaptic successes and failures. This is problematic given that accurate classification of small synaptic events relies on appropriate discrimination from noise levels that are inherently different between hyperpolarized and depolarized states. The generally unknown and variable presence of rectifying GluA2-lacking AMPA-Rs (that differentially contribute to the EPSC at depolarized and hyperpolarized states) may also contribute to this classification ambiguity. Our method works on the failure rates directly, and therefore it assumes that there are no systematic biases inherent in this signal classification. Examining and correcting such classification biases from the raw analog signals is beyond the scope of our MLE method, but future work is needed to address these concerns.

Our power analysis additionally provides key insights for more modern binary synapse classification tools, including 2-photon glutamate uncaging (29). Our results demonstrate that in these cases, a dramatic boost to statistical power (∼3-fold) can be obtained through constrained hypothesis testing using analytical tools – for example, testing against the null hypothesis that there are no silent synapses by employing log-likelihood ratio tests. We suggest that such hypothesis-based testing constitutes a viable alternative to classical Chi-squared statistical tests when using modern binary classification techniques.

The FRA formalism relies on several rigid, core assumptions. First, FRA assumes that the release probability distributions for silent and nonsilent synapses are equivalent, since this is a core simplification needed to derive the FRA equation (see STAR Methods). However, it is entirely possible that silent synapses may have distinct release probabilities (51), which might bias the estimates. Second, FRA assumes that NMDA opening is the sole parameter that changes between recordings at hyperpolarized and depolarized potentials, and that other parameters such as synaptic release probability are constant at both holding potentials. However, several synapses have been shown to be regulated by depolarization-induced release of retrograde messengers, which can either increase (42) or decrease (43-44) release probability. In these circuits, the FRA approach may be fundamentally inadmissible, as the returned computation would reflect the effects of the retrograde messengers rather than the silent synapse fraction. Contrary to the FRA approach, synthetic likelihoods can incorporate the expected action of retrograde messengers. By comparing the likelihood with and without incorporating these changes in release probability, standard model validation methodologies (38) can assess whether the changes in failures rates truly reflect a change in silent synapse fraction. Altogether, the formalism developed herein provides not only an improved analytical tool to estimate silent synapse fraction from electrophysiological recordings, but also an independent means to assess assumption violations.

## Author Contributions

Conceptualization, M.B.L., K.F.H.L, C.S, R.N, J.-C.B; Methodology, M.B.L.; Software, M.B.L.; Formal Analysis, M.B.L.; Investigation, M.B.L.; Writing - Original Draft, M.B.L.; Writing - Review & Editing, M.B.L, R.N and J.-C.B.; Supervision, J.-C.B and R.N; Funding Acquisition, J.-C.B.

## STAR Methods

### RESOURCE AVAILABILITY

#### LEAD CONTACT

Further information and requests for resources and reagents should be directed to and will be fulfilled by the Lead Contact, Michael Lynn.

#### MATERIALS AVAILABILITY

This study did not generate new unique reagents.

#### DATA AND CODE AVAILABILITY

The code generated during this study is available at https://github.com/micllynn/SilentMLE.

### EXPERIMENTAL MODEL AND SUBJECT DETAILS

#### MICE

This study employed male mice (C57BL/6; P14-P15; 10-15g). Mice were group housed before experimental procedures. All experiments and procedures were performed in accordance with approved procedures and guidelines set forth by the University of Ottawa Animal Care and Veterinary Services.

### METHOD DETAILS

#### DERIVATION OF FRA EQUATION

Let *n*_*s*_ represent the number of silent synapses in some ensemble, *n*_*a*_ the number of active synapses, *F*_*h*_ the failure rate at hyperpolarized potentials (*n*_*a*_ synapses active) and *F*_*d*_ the failure rate at depolarized potentials (*n*_*a*_ + *n*_*s*_ synapses active since NMDA-receptor-containing silent synapses can only conduct under depolarized settings). Then:

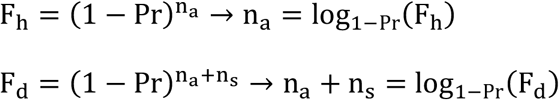

The silent fraction is 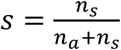, so:

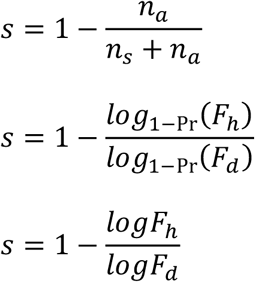

This can be written as a percentage:

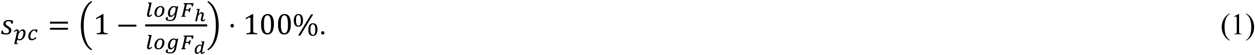

#### SIMULATIONS WITH ACTIVE SYNAPSES ONLY

We simulated single active (nonsilent) synapses or groups of active synapses. For each synapse *i* we assigned a release probability by drawing from a uniform distribution *Pr*_*i*_ ∼ *U*(0,1).

We numerically stimulated stochastic neurotransmitter release at hyperpolarized voltages (*V*_*h*_; only active synapses) and depolarized voltages (*V*_*d*_; active and silent synapses). 50 trials were simulated at each of *V*_*h*_ and *V*_*d*_. Trials were modelled as Bernoulli trials with failure rate 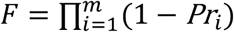 over *m* synapses considered, where 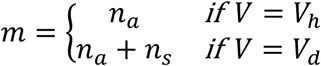. This yielded 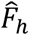 and 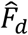, which are estimates of the true failure rate over 50 trials at *V*_*h*_ and *V*_*d*_ respectively. The FRA calculation was performed as reported in equation (1), with 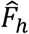 and 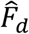 as the equation inputs. Simulated currents (for purposes of illustration only) were generated by modeling the total current as a sum of all successfully releasing synapses, where each success is treated as generating a stochastic unitary current with an (arbitrary) mean amplitude of 10pA. However, all simulations proceeded using idealized Bernouilli trials to compute the stochastic failure rates (i.e. no mis-classification of successes and failures)

#### SIMULATIONS WITH ACTIVE AND SILENT SYNAPSES

In this section, we describe our simulations of minimum-simulation electrophysiological experiments, where a larger set of synapses with defined release probabilities and some fraction silent are iteratively reduced into some experimentally feasible subset on which to perform experiments. This iterative reduction models a gradual reduction in electrical stimulation intensity that an electrophysiologist would perform.

Assume the true fraction of silent synapses in a population is *s*_*t*_. For each replicate, an initial set of silent and active synapse numbers {*n*_*s*_, *n*_*a*_} were each chosen from a binomial distribution with an initial population size of *n*_*t*_ = 100 synapses and *s*_*t*_ silent probability (e.g. the distribution of *n*_*s*_ silent synapses was 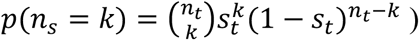. Release probabilities for each synapse were I.I.D and were drawn from either a uniform or gamma distribution as noted in the text, producing a vector of release probabilities 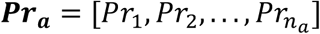 for active synapses and 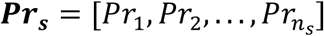 for silent synapses. The synapse set {*n*_*s*_, *n*_*a*_} and associated release probability set {***Pr***_***s***_, ***Pr***_***a***_} were then subjected to iterative rounds of reduction, which yielded an experimentally feasible subset.

Iterative reduction of the synapse sets proceeded until the failure rate criterion was met: 0.2 < *F*_−70_ < 0.8. Here, 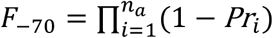 where *Pr*_*i*_ ∈ ***Pr***_***a***_. F is analytically calculated at each iteration, rather than estimated using Monte Carlo simulations, for purposes of computational efficiency. Iterative reduction consisted of the following sequential steps: 1) Using *p*_*elim*_ = 0.1 as the elimination probability, stochastically eliminate *p*_*elim*_(*n*_*s*_ + *n*_*a*_) synapses along with their associated release probabilities in {***Pr***_***s***_, ***Pr***_***a***_}. 2) After each elimination, calculate *F*_−70_. 3). If *F*_−70_ falls within criterion range, then store {*n*_*s*_, *n*_*a*_} and {***Pr***_***s***_, ***Pr***_***a***_}. If 1_−70_ does not fall within criterion range, loop back to 1) and repeat.

Together, this iterative decrease yielded Monte Carlo-simulated subsamples (replicates) of active and silent synapses along with associated release probabilities {***Pr***_***s***_, ***Pr***_***a***_}. For each replicate, we used {***Pr***_***s***_, ***Pr***_***a***_} to simulate 50 Bernoulli trials at each of *V*_*m*_ = −70*mV* (*n*_*a*_ synapses) and *V*_*m*_ = +40*mV* (*n*_*a*_ + *n*_*s*_ synapses) similar to above, yielding stochastic failure rate estimates, *F*_−70_ and *F*_+40_. These estimates were used as input to the FRA estimator, to return one silent fraction estimate, *s*_*FRA*_, per Monte Carlo simulation.

For each ground truth silent value, *s*_*t*_, we obtained ∼20 000 replicates which each yielded an estimate, *s*_*FRA*_. These numerically simulated distributions formed the basis of our bias and variance analysis the FRA estimator. For each *s*_*t*_, we can then compute:

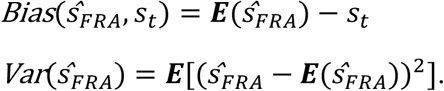

### EXPERIMENTAL MEASUREMENTS OF FRA

Hippocampal slices (300um thick) were made from P14-15 wild-type (C67/Blk) mice using a previously described technique (36). In brief, mice were anesthetized using isofluorane, decapitated, and the brain was extracted and placed in ice-cold choline-based cutting solution for the duration of the cut. Cutting solution was comprised of 119 mM choline-Cl, 2.5 mM KCl, 1 mM CaCl2, 4.3 mM MgSO4-7H2O, 1 mM NaH2PO4, 1.3 mM sodium L-ascorbate, 26.2 mM NaHCO3, and 11 mM glucose, and was equilibrated with 95% O2 and 5% CO2. 300uM coronal sections were made of the hippocampus. Slices were recovered in a chamber containing standard Ringer’s solution of the following composition: 119 mM NaCl, 2.5 mM CaCl2, 1.3 mM MgSO4-7H2O, 1 mM NaH2PO4, 26.2 mM NaHCO3, and 11 mM glucose, at a temperature of 37 °C, continuously bubbled with a mixture of 95% O2 and 5% CO2. Slices were recovered for 1 h in the recovery chamber and equilibrated to a temperature of ∼25 °C until recordings were performed.

Neurons were visualized in the CA1 subfield using an upright microscope (Examiner D1 equipped with differential interference contrast (DIC) (Zeiss; 40×/0.75 N.A. objective). Using 4-6mOhm fire-polished glass electrodes (Sutter Corporation) pulled on a on a Narishige PC-10 pipette puller, CA1 subfield neurons were recorded in the whole-cell patch-clamp configuration at room temperature (20-22C) with a cesium-based internal comprised of 115 mM cesium methane-sulfonate, 5 mM tetraethylammonium-Cl, 10 mM sodium phosphocreatine, 20 mM Hepes, 2.8 mM NaCl, 5 mM QX-314, 0.4 mM EGTA, 3 mM ATP (Mg2+), and 0.5 mM GTP (pH 7.25) (adjusted with CsOH; osmolarity, 280–290 mOsmol/L). As an external solution, Ringer’s solution was continuously circulated. Whole-cell recordings were carried out using an Axon Multiclamp 700B amplifier, sampled at 10 kHz, digitized with an Axon Digidata 1440A (or 1550) digitizer, and filtered at 2 kHz. An electrical stimulation electrode was placed apposed to the apical dendritic arbor to activate Schaffer collaterals. Stimulation intensity was tuned such that ∼50% synaptic failures and successes each were recorded.

The FRA protocol consisted of 100 sweeps of electrical stimulation (0.1ms pulse width; 10-15 seconds sweep-to-sweep time). 50 sweeps were performed at *V*_*m*_ = −70*mV* and 50 sweeps were performed at *V*_*m*_ = +40*mV*. This FRA protocol was performed in triplicate on single neurons, with FRA iterations immediately following one another. Recordings were discarded when the access resistance changed by >30%. Liquid junction potential was not compensated for. The data was analyzed offline with the aid of a custom Python module written by the author and available online at: https://github.com/micllynn/synappy. Synaptic failures and successes were determined in an automated manner by setting a threshold for the baseline-subtracted electrically evoked current to exceed to be considered a success. The FRA calculation was performed as indicated in equation (1), by comparing the failure rate at -70mV and +40mV. Mean estimate spread was calculated by subtracting the minimum FRA-estimated silent synapse fraction from the maximum FRA-estimated silent synapse fraction within the same cell.

All experiments and procedures were performed in accordance with approved procedures and guidelines set forth by the University of Ottawa Animal Care and Veterinary Services.

#### THE IMPROVED MAXIMUM-LIKELIHOOD ESTIMATOR

For a jointly observed pair of failure rates at hyperpolarized and depolarized potentials, 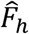 and 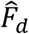, we wish to estimate the most likely *s*_*t*_. If we had an analytically tractable likelihood function, we could use likelihood-based inference techniques (e.g. maximum likelihood estimation) to find our most likely *s*_*t*_. However, two factors make it difficult to analytically derive a likelihood function: first, the number of active and silent synapses in each sampled subset are only probabilistically related to *s*_*t*_; and second, each synapse has release probabilities Pr which are a complex function of the starting population’s Pr and a bias imposed by the number of other (active) synapses present in that particular subset.

To circumvent these obstacles, we use a statistical inference approach inspired by recent treatments of noisy dynamical systems (33). While each individual simulation of synaptic release trains contains a large amount of information, we can extract summary statistics for each simulation and construct a synthetic likelihood function yielding the probability of obtaining these summary statistics at each underlying value of *s*_*t*_. We choose the biased estimator *ŝ*_*FRA*_ as the summary statistic as it contains information about both 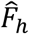 and 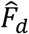 (see below for a detailed discussion about this choice of summary statistic). Our synthetic likelihood function can therefore be written in terms of the summary statistic: 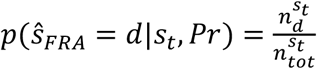, where *d* is a given value of observed data, 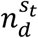 is the count of Monte Carlo-simulated replicates using the parameters *s*_*t*_, *Pr* which satisfy *ŝ*_*FRA*_ = *d* (i.e. the observation is obtained), and 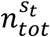 is the total number of Monte Carlo-simulated replicates using the parameters *s*_*t*_, *Pr*. As above, *s*_*t*_ refers to the true silent fraction and *Pr* denotes a synaptic release probability distribution.

When more than one piece of data is acquired, is convenient to write a joint log-likelihood function over a given vector of observations ***D*** = (*d*_1_, *d*_2_, …, *d*_*n*_). We can express this as 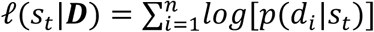. We can then construct an MLE estimator for *s*_*t*_ given some vector of measured data &, expressed as 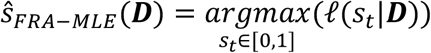. This estimator’s bias and variance was evaluated in an identical way as the FRA estimator (above), and the MLE estimator was found to exhibit no bias and significantly lower variance.

Due to the stochastic and finite nature of the simulations needed to generate *ℓ*(*s*_*t*_|***D***), and to save computational resources, we have smoothed the numerical estimate for *ℓ*(*s*_*t*_|***D***) using a Savitsky-Golay filter (window length = 0.1 silent fraction; polynomial order = 3) before computing 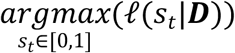.

Two caveats to this approach: First, note that instead of constructing *p*(*ŝ*_*FRA*_|*s*_*t*_) point-by-point with some step size *δs*_*t*_ → 0, we could conceivably model the likelihood function as a continuous probability density over all values of *s*_*t*_, by estimating parameters of a normal distribution similar to (33). Here, one could compute 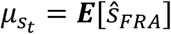 and 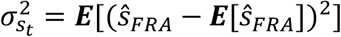 for each *s*_*t*_. However, we have established that the distribution of *ŝ*_*FRA*_ for each *s*_*t*_ has nonzero skewness and kurtosis, making it ineligible for normal distribution fitting.

Second, while we chose one summary statistic *ŝ*_*FRA*_ to construct a synthetic likelihood function, it is also possible to directly use the failure rates 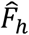 and 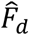 as two summary statistics. This would appear to be advantageous because we are using two summary statistics instead of one. However, this does not provide better estimation capabilities because the distributions for 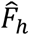 show no relationship with *s*_*t*_ in our simulations. Therefore, we chose a single summary statistic *ŝ*_*FRA*_.

It is also formally possible that some summary statistic extracted from the raw event traces themselves (such as the variance of amplitudes of successful trials) could provide more information about the fraction of silent synapses in some ensemble than the failure rates alone. However, the unitary event amplitudes of AMPA-R and NMDA-R responses are typically not equivalent, nor are their standard deviations. Furthermore, there is typically more noise in recordings when a neuron is clamped at a depolarized potential. Since all of these factors can vary between experimental setups and brain regions, it is not trivial to provide a generalizable solution based on summary statistics of raw event traces. For these reasons, we took a failure-rate-based approach for the summary statistic.

### POWER ANALYSIS OF ESTIMATORS FOR *s*_*t*_

We employed a power analysis approach to compare the statistical power of different techniques for silent synapse detection. Parameters for power analyses were *α* = 0.05 (false positive rate) and *β* = 0.2 (false negative rate) unless otherwise noted.

Given two populations of synapses, *P*_1_ and *P*_2_, having distinct silent fractions 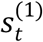 and 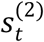 silent fractions, respectively. We want to determine *n*_*min*_, the minimum number of samples required to discriminate these populations with some estimation technique. To do this, we wish to express statistical power, *p*, as a function of the number of samples: *p*(*n*). We can then find an *n*_*min*_ such that *p*(*n* > *n*_*min*_) < *β*. Our approach to explore *p*(*n*) was to rely on Monte Carlo sampling methods paired with an efficient binary search algorithm to search the space of possible *n* ∈ [0,2^11^] (an arbitrary maximum limit on *s* to search, due to computational constraints). Here, at each iteration, a sample size guess *n*_*guess*_ is made. After Monte Carlo simulations and an evaluation of statistical power (whether *p*(*n*_*guess*_) < *β* or *p*(*n*_*guess*_) > *β*). *n*_*guess*_ is adjusted up (if insufficient statistical power), or down (if sufficient statistical power). The amount that *n*_*guess*_ is adjusted by is halved at each time point in order to converge on the true value for *n*_*min*_ in precisely *t* = 11 iterations (i.e. the algorithm has order *O*(*log*_2_(*n*)) where n is the initial guess.) The algorithm is described in the Python source code. Briefly:

0) Using the Monte Carlo methodology described above, generate numerical probability distributions for estimates of 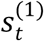 and 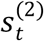. These distributions are denoted ***p***^(**1**)^(***ŝ***_***FRA***_) and ***p***^(**2**)^(***ŝ***_***FRA***_).
1) Set *n*_*guess*_ = 2^11^ = 2048. Draw *k* = 10000 independent replicates, each composed of *n*_*guess*_ paired experimental samples sampled from ***p***^(**1**)^(***s***_***FRA***_) and ***p***^(**2**)^(***s***_***FRA***_). This yields corresponding replicate matrices ***D***_1_ and ***D***_2_, which have dimension (*k, n*_*guess*_).
2) *Evaluation of statistical power*. For each one of the *k* columns in ***D***_1_ and ***D***_2_, use a Wilcoxon Rank-Sum statistical test and record whether the replicate is classified as a positive with *p* < *α*. Since the populations have distinct silent fractions, all positives are true positives so we can record the number of true positive *n*_*TP*_. Compute the false-negative rate as *FNR* = 1 − *n*_*TP*_/*k*.
3a) If *FNR* < *β*: Update 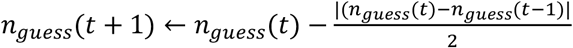. Loop back to (1).
3b) If *FNR* > *β*: Update 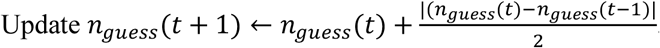. Loop back to (1).
4) Repeat until |*n*_*guess*_(*t*) − *n*_*guess*_(*t* − 1)| = 1. Then *n*_*min*_ = *min*{*n*_*guess*_(*t*), *n*_*guess*_(*t* − 1)}.

In Fig. 4C, we test each of the four estimators (*fra, binary, fra-mle, binary(llr)*) by testing the null hypothesis that *P*_1_ is silent. So we fix 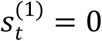 while 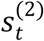 ∈ [0,1) and compute 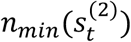 for each estimator:

i. For *ŝ*_*FRA*_(*fra*) the computation of 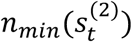 proceeds numerically as described in the algorithm above.
ii. For the binary sampling case (*binary*), the algorithm was identical to above, except instead of drawing estimates from a Monte Carlo-simulated distribution, we simply considered each draw of *n*_*guess*_ samples as constituting Bernoulli trials with 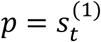 or 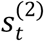. In step 2, a Chi-Squared test was used instead of a Rank-Sum test as the data is categorical (silent or active binarization).
iii. For *ŝ*_*FRA*−*MLE*_ (*fra-mle*), we employ a log-likelihood (LLR) approach. We first calculate likelihood functions as described above. In all cases the null hypothesis is *H*_0_: *s*_*t*_ = 0 and the alternate hypothesis is *H*_*A*_: *s*_*t*_ *= ŝ*_*FRA*−*MLE*_. We apply Wilks’ theorem, calculating a test statistic *D* = −2[*ℓ*(*s*_*t*_ = *ŝ*_*FRA*−*MLE*_) − *ℓ*(*s*_*t*_ = 0)] where *D* ∼ *χ*^2^(*df* = 1). This allows us to employ a Chi-Squared test on D in place of a Wilcoxon Rank-Sum test (step 2 of the algorithm above). All other components of the algorithm are identical.
iv. For a binary sampler incorporating a model (*binary(llr)*), we derive an analytical expression for minimum sample size, *n*_*min*_, with a LLR testing approach (starting from the binomial distribution): 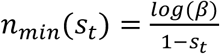.

### QUANTIFICATION AND STATISTICAL ANALYSIS

All numerical simulations and statistical analysis were performed using custom scripts employing standard functions of technical computing libraries NumPy (39) and SciPy (40) in a Python 3.6 environment. Further details about statistical tests are described in the Figure captions, Results and Methods of each respective section. In all cases, significance was defined as p < 0.05.

### KEY RESOURCES TABLE

| REAGENT or RESOURCE | SOURCE | IDENTIFIER |
| --- | --- | --- |
| Experimental Models: Organisms/Strains |  |  |
| Mouse: C57BL/6 | Charles River Labs | C57BL/6NCrl |
| Software and Algorithms |  |  |
| Python | Python Software Foundation | Python 3.6 |
| NumPy | NumPy | NumPy 1.18.1 |
| SciPy | SciPy | SciPy 1.4.1 |
| SilentMLE Python Package | This paper | https://github.com/micllynn/silentmle |

