## Supplemental Figures for "A synthetic likelihood solution to the silent synapse estimation problem"

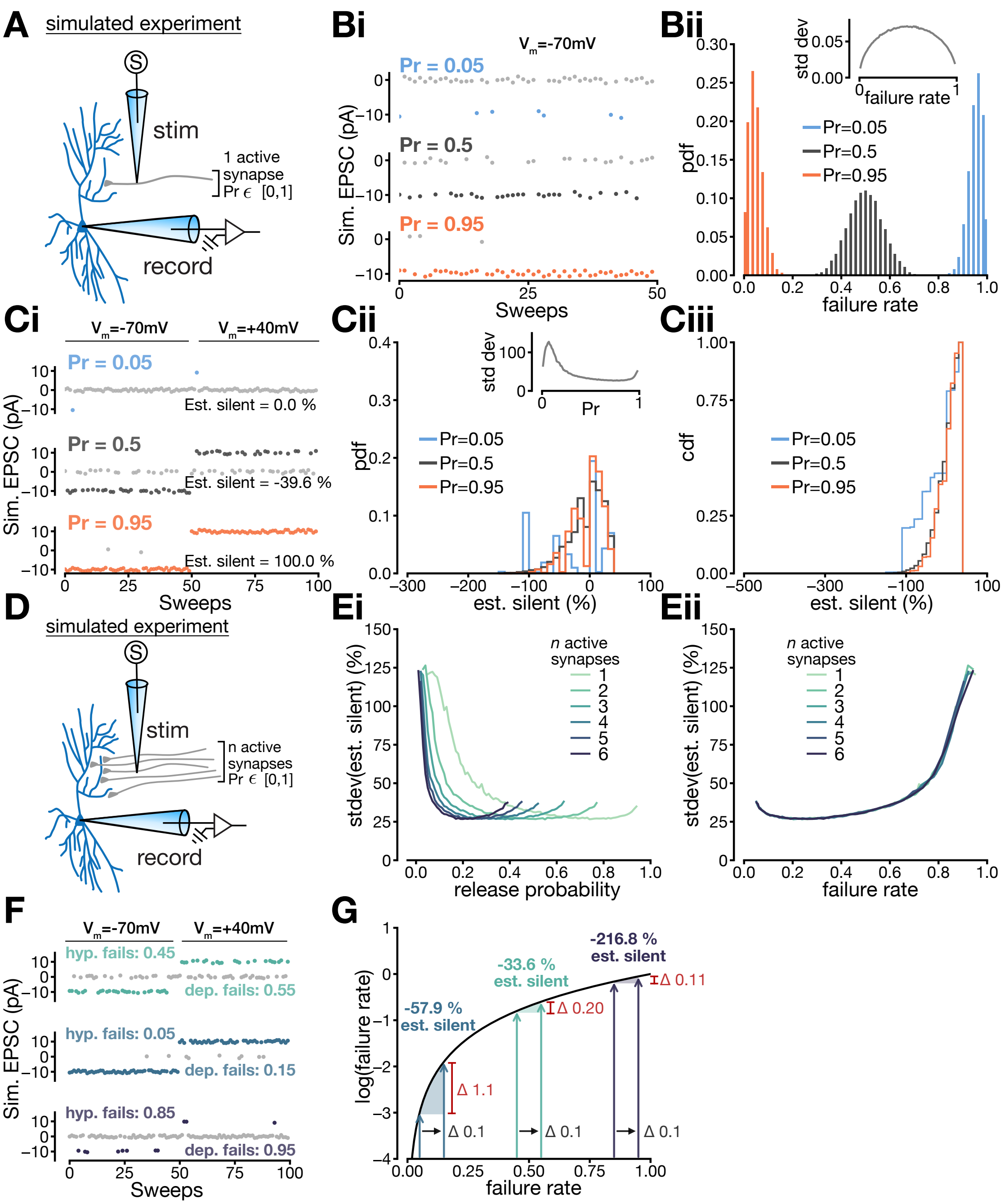


**Supplemental Figure 1**: The poor estimation characteristics of the FRA equation do not vary significantly over the parameter space of release probabilities and synapse numbers. **A:** Depiction of simulation with one active synapse. **Bi:** Representative simulations for failures and successes at Vm = -70mV for Pr = 0.05, Pr = 0.5 and Pr = 0.95. **Bii:** Representative distributions for failure rate at Pr = 0.05, Pr = 0.5 and Pr = 0.95. *Inset*; failure rate distributions have standard deviations which are parabolically related to release probability. **Ci:** Representative simulations and silent fraction estimates for full FRA experiments with Pr = 0.05, Pr = 0.5 and Pr = 0.95. **Cii:** Representative distributions for estimated silent fraction at Pr = 0.05, Pr = 0.5 and Pr = 0.95. *Inset*; in contrast to failure rate, the distributions for estimated silent fraction exhibit similar standard deviations across the range of possible release probability values. **Ciii:** Same as **Cii**, but the distributions are plotted as cumulative probability densities.

**D:** Depiction of simulation with multiple active synapses. **Ei:** Standard deviation of the estimated silent fraction as a function of release probability and number of active synapses. **Eii:** Standard deviation of the estimated silent fraction as a function of failure rate and number of active synapses. **F:** Representative traces of three simulated FRA experiments where the hyperpolarized and depolarized failure rates differ by 10%. **G:** Depiction of the FRA estimates in F, showing log(failure rate) plotted against failure rate. An increase in failure rate of 0.1, at higher absolute failure rates, produces correspondingly smaller effects in log(failure rate).

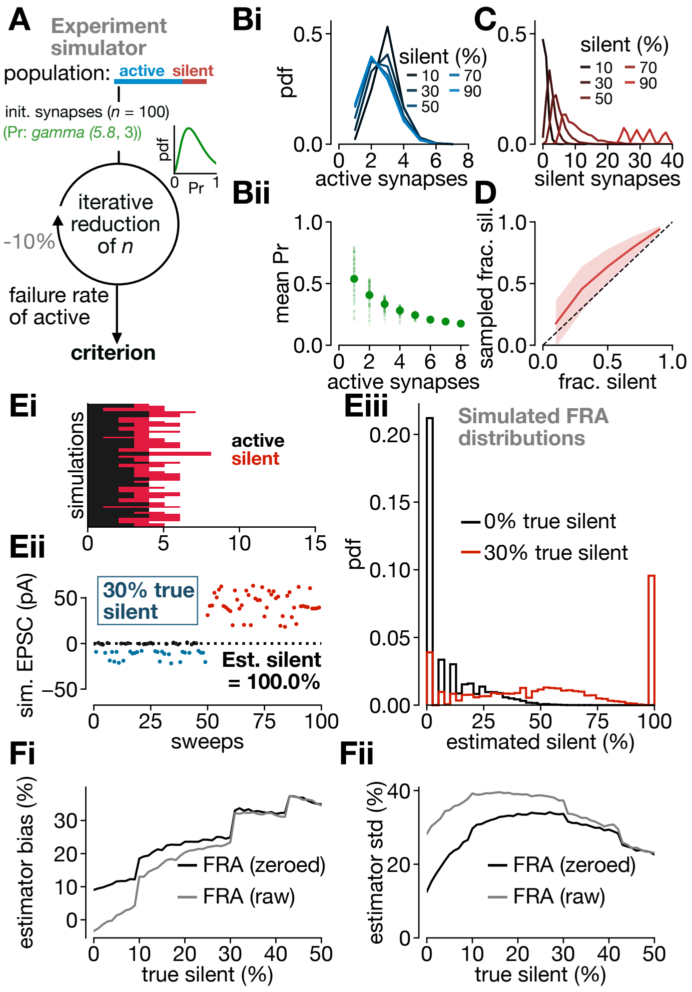


<u>**Supplemental Figure 2**</u>: Experimentally bounded simulations with a gamma release probability distribution (23). **A:** Schematic depicting experimental simulator approach. **Bi:** Probability distribution of number of active synapses obtained during each simulation. **Bii:** Release probability of active synapses. Small dots denote release probability for each simulation, while large dots denote the mean across simulations. **C:** Probability distribution of number of silent synapses obtained during each simulation. **D:** Sampled fraction silent is shown against the true fraction silent. **Ei**: Depiction of returned samples of silent/active synapses. **Eii**: Depiction of a simulated FRA experiment performed on a sample of synapses returned from the iterative reduction. **Eiii**: Distribution of FRA estimates for two true silent values. **Fi:** FRA estimator bias for non-zeroed and zeroed distributions. **Fii**: FRA estimator standard deviation for non-zeroed and zeroed distributions.

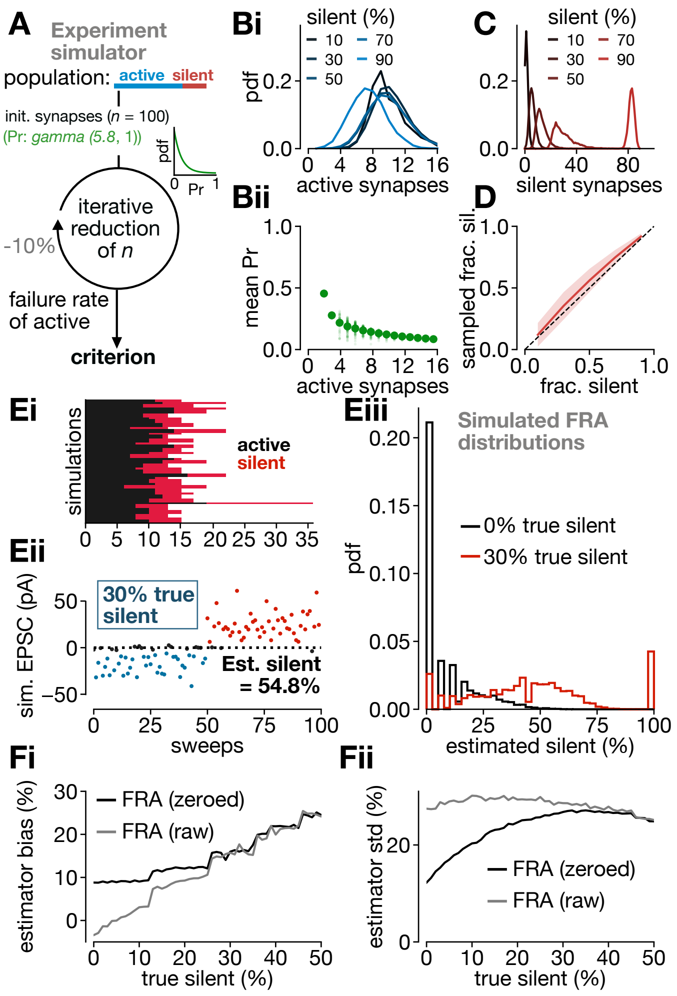


**<u>Supplemental Figure 3</u>**: Experimentally bounded simulations with a gamma release probability distribution biased towards low values (). **A:** Schematic depicting experimental simulator approach. **Bi:** Probability distribution of number of active synapses obtained during each simulation. **Bii:** Release probability of active synapses. Small dots denote release probability for each simulation, while large dots denote the mean across simulations. **C:** Probability distribution of number of silent synapses obtained during each simulation. **D:** Sampled fraction silent is shown against the true fraction silent. **Ei**: Depiction of returned samples of silent/active synapses. **Eii**: Depiction of a simulated FRA experiment performed on a sample of synapses returned from the iterative reduction. **Eiii**: Distribution of FRA estimates for two true silent values. **Fi:** FRA estimator bias for non-zeroed and zeroed distributions. **Fii**: FRA estimator standard deviation for non-zeroed and zeroed distributions.

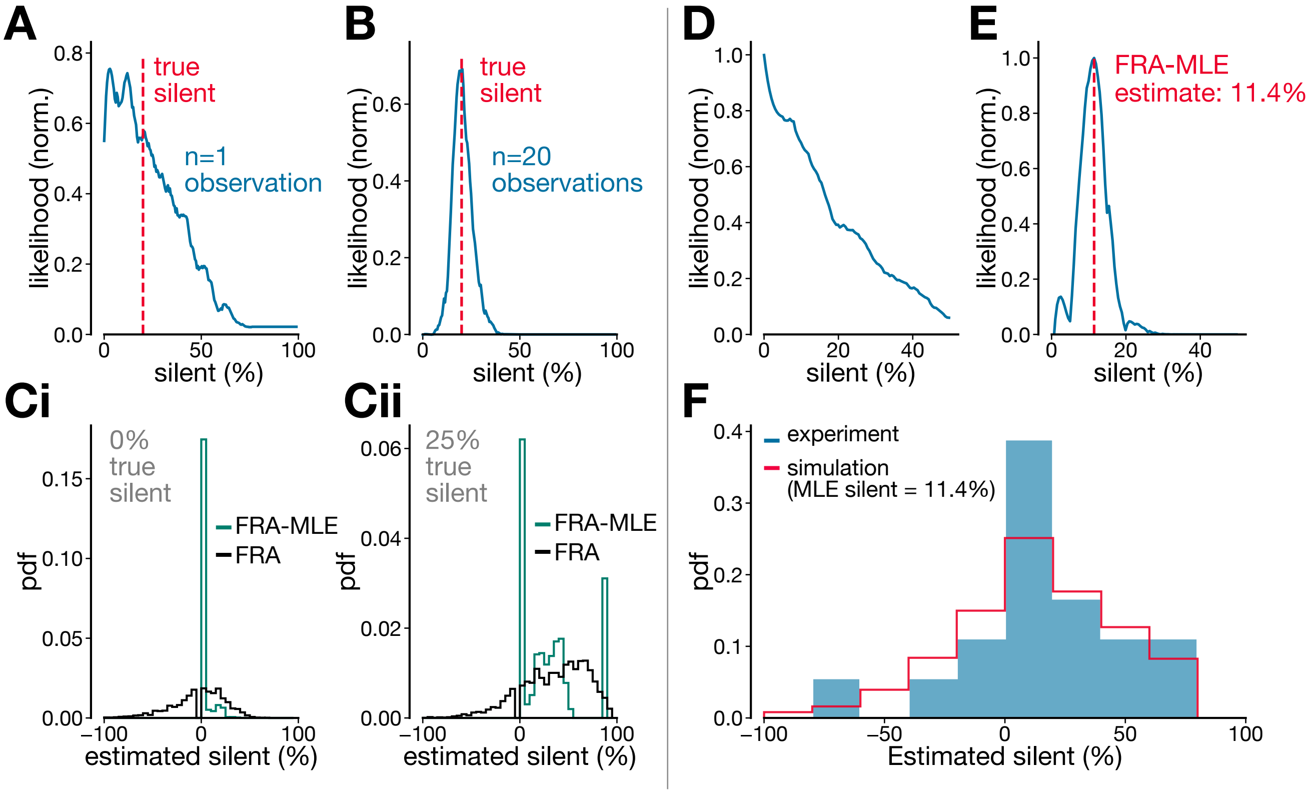


**<u>Supplemental Figure 4</u>**: A maximum likelihood estimator for silent synapse fractions. **A-C** depict simulations verifying the performance of the FRA-MLE estimator. **A**: Given a single observation from an FRA distribution, FRA-MLE returns likelihood function which shows that multiple silent fractions are consistent with this one observation. **B:** When 20 observations from the same FRA distribution are considered together by FRA-MLE as a joint likelihood function, a highly precise estimate of the true silent fraction is possible. **C**: Given a true silent fraction of 0% (**Ci**) or 25% (**Cii**), FRA-MLE on single observations returns low-variance, precise estimates, while FRA returns high-variance, imprecise estimates. **D-F** depict an application of the FRA-MLE estimator to experimental data in hippocampus at the CA3-CA1 synapse (full data shown in Fig. 2). **D:** Likelihood function returned by FRA-MLE from a single experimental observation. **E:** The joint likelihood function, across all experimental observations, returned by FRA-MLE. The FRA-MLE estimate from this joint likelihood function is 11.4% silent. **F:** A comparison of the distribution of FRA-estimated silent values (blue) overlaid with a simulated distribution of estimated silent values, calculated at the maximum likelihood estimate from FRA-MLE (11.4% silent). The distributions are not statistically different ($p = 0.99$; KS-test, two-tailed)
